# Arabidopsis pollen Prolyl-hydroxylases P4H4/6 are required for correct hydroxylation and secretion of LRX11 in pollen tubes

**DOI:** 10.1101/2022.11.16.516804

**Authors:** Ana R. Sede, Diego L. Wengier, Cecilia Borassi, Martiniano Ricardi, Sofía C. Somoza, Rafael Aguiló, José M. Estevez, Jorge P. Muschietti

**Affiliations:** Instituto de Investigaciones en Ingeniería Genética y Biología Molecular, Dr. Héctor Torres (INGEBI-CONICET), Vuelta de Obligado 2490, Buenos Aires, C1428ADN, Argentina; Fundación Instituto Leloir and IIBBA-CONICET, Av. Patricias Argentinas 435, Buenos Aires, Argentina; Instituto de Fisiología, Biología Molecular y Neurociencias (IFIByNE-CONICET), Universidad de Buenos Aires, Buenos Aires, Intendente Güiraldes 2160, Ciudad Universitaria, C1428EGA, Buenos Aires, Argentina; Centro de Biotecnología Vegetal (CBV), Facultad de Ciencias de la Vida, Universidad Andrés Bello and ANID - Millennium Nucleus for the Development of Super Adaptable Plants (MN-SAP), Santiago, Chile; Departamento de Biodiversidad y Biología Experimental, Facultad de Ciencias Exactas y Naturales, Universidad de Buenos Aires, Intendente Güiraldes 2160, Ciudad Universitaria, Pabellón II, C1428EGA, Buenos Aires, Argentina; Institut de Biologie Moléculaire des Plantes, CNRS, Université de Strasbourg, 67000 Strasbourg, France; Department of Chemical Engineering, Stanford University & HHMI Shriram Center, Stanford, CA, USA; Laboratory of Biochemistry, Wageningen University, Stippeneng 4, Wageningen, The Netherlands; Department of Biology, University of Padova, 35131, Padova, Italy

**Keywords:** *Arabidopsis thaliana*, glycoprotein secretion, LRXs, P4Hs, pollen germination, pollen tubes, plant cell wall

## Abstract

Major constituents of the plant cell walls are structural proteins that belong to the Hydroxyproline-rich glycoprotein family. Leucine-rich repeat extensis are contain a leucine-rich domain and a C-terminal domain with repetitive Ser-Pro(3-5) motifs plausible to be glycosylated. We have demonstrated that pollen-specific LRX8-11 from *Arabidopsis thaliana* are necessary to maintain the integrity of the pollen tube cell wall during polarized growth. In classical EXTs and likely in LRXs, proline residues are converted to hydroxyproline by Prolyl-4-hydroxylases, thus defining novel *O*-glycosylation sites. In this context, we aimed to determine whether hydroxylation and subsequent *O*-glycosylation of Arabidopsis pollen LRXs are necessary for their proper function and cell wall localization in pollen tubes. We hypothesized that pollen-expressed P4H4 and P4H6 catalyze the hydroxylation of the proline units present in Ser-Pro(3-5) motifs of LRX8-LRX11. Here, we show the *p4h4-1 p4h6-1* double mutant exhibits a significant reduction in pollen germination rates and a slight reduction in pollen tube length. Pollen germination is also inhibited by specific P4Hs inhibitors, suggesting that prolyl hydroxylation is required for pollen tube development. Plants expressing *pLRX11::LRX11-GFP* in the *p4h4-1 p4h6-1* background show partial relocalization of LRX11-GFP from the pollen tube tip apoplast to the cytoplasm. Finally, IP-MS- MS analysis revealed a decrease in oxidized prolines in LRX11-GFP in the *p4h4-1 p4h6-1* background when compared to *lrx11* plants expressing *pLRX11::LRX11-GFP*. Together, these results suggest that P4H4 and P4H6 are required for pollen germination and are also involved in LRX11 hydroxylation necessary for its localization at the cell wall of pollen tubes.

**One Sentence Summary:** Pollen-expressed P4H4 and P4H6 are required for pollen germination and for proper hydroxylation and secretion of LRX11 in pollen tubes.

## Introduction

Plant cell walls are mainly composed of structural proteins that belong to the Hydroxyproline-rich glycoprotein (HRGP) superfamily that are embedded in a polysaccharide matrix contributing in cell wall architecture. Members of HRGPs are subdivided according to their glycosylation pattern in: hyperglycosylated arabinogalactan proteins (AGPs), moderately glycosylated extensins (EXTs) and slightly glycosylated proline-rich proteins (PRPs) (Showalter *et al*., 2010). Glycosylation is crucial for proper folding and biological function of plasma membrane and cell wall proteins. Plant specific *O*- glycosylation predominantly consists of the attachment of *O*-linked glycans, mainly linear arabinosyl units in EXTs, to the hydroxyl group of hydroxyproline (Hyp), abundant in HRGPs (Strasser, 2016; Marzol *et al*., 2018). Proline (Pro) residues present in characteristic repetitive Ser-Pro(3-5) motifs of EXTs undergo hydroxylation followed by *O*-glycosylation in the ER-Golgi apparatus, and proper folded proteins are secreted in the cell wall where participate in the formation of a covalent network mediated by Tyr cross-linking (Hijazi *et al*., 2014). In this context, hydroxylation, as the first post translational modification necessary to define novel *O*-glycosylation sites, is catalyzed by members of the 2- oxoglutarate-dependent dioxygenases family known as Prolyl-4-hydroxylases (P4Hs) (Gorres and Raines, 2010). In root hairs, P4H2, P4H5 and P4H13 are required for a correct cell wall assembly and hence for polarized growth of Arabidopsis root hairs through peptidyl-proline hydroxylation of EXTs (Velasquez *et al*., 2011, 2015a,b). Interestingly, an enzymatic *in vitro* competition assay demonstrated that P4H5 hydroxylate preferentially polyproline present in EXT-type substrates rather than AGPs-type (Velasquez *et al*., 2015b). Once proline-hydroxylation is complete, a stepwise elongation of sugar chains takes place and this process involves different groups of glycosyltransferases (GTs; reviewed in Showalter and Basu, 2016). Hydroxyprolil-*O*-arabinosyl-transferases (HPATs) have been identified as the enzymes responsible for the attachment of the first arabinose unit to the Hyp (Ogawa-Ohnishi and Matsubayashi, 2015). It was shown that HPATs activity and hence arabinosylation are necessary for the maintenance of tip growing cells both in *Arabidopsis thaliana* and the moss *Physcomitrella patens* (MacAlister *et al*., 2016). Furthermore, Arabidopsis *hpat1/2/3* triple knockout mutants displayed strong male sterility due to morphological defects in pollen tube elongation (MacAlister *et al*., 2016).

Leucine-rich repeat EXTs (LRXs) are extracellular proteins of the EXTs family, crucial to sustain polarized growth of root hairs (Baumberger, Ringli and Keller, 2001) and pollen tubes (Baumberger *et al*., 2003; Ndinyanka Fabrice *et al*., 2017; Sede *et al*., 2018; Wang *et al*., 2018). Pollen-specific LRX8-11 interact by their N-terminal domain with the Rapid Alkalization Factor (RALF) peptides (RALF4/19) at the cell wall to activate the ANXUR signaling pathway involved in maintaining proper polarized growth (reviewed in Somoza *et al*., 2020). It was shown that the extensin domain of LRX1 is essential for protein function but its insolubilization at the root hair cell wall is independent of the presence of Tyr residues, indicating the existence of other strong interactions through its EXT domain (Ringli, 2010). These interactions might involve the presence of glycosylation, since LRXs contain an EXT-like C-terminal domain with repetitive motifs of Ser-Pro(3-5) plausible to be glycosylated (Sede *et al*., 2018). A previous study identified LRX3, from the vegetative clade, as a P4H5 interactor in a yeast two-hybrid assay (Velasquez *et al*., 2011). However, the role of P4Hs and therefore of the hydroxylation of pollen-specific EXTs during pollen germination and pollen tube polarized growth remain unknown. In this context, we identified and characterized two pollen-expressed P4Hs and to evaluate the importance of proline-hydroxylation and subsequent *O*-glycosylation for a proper EXTs cell wall assembly. Here, we determined by several approaches that pollen-specific LRXs are putative targets of P4Hs enzymes and that proline hydroxylation is necessary for proper LRXs secretion and their anchoring at the cell wall.

## Results

### Identification of pollen-expressed P4H4 and P4H6

The Arabidopsis thaliana genome contains 13 loci that code for *P4Hs*. According to the public databases (Affymetrix ATH1 microarray data available from https://genevestigator.com; Loraine *et al*., 2013), two *P4H* genes, *P4H4* and *P4H6*, are highly expressed in mature pollen (**Supplemental Figure 1**). Full length protein sequence alignment and phylogenetic analysis of Arabidopsis P4Hs revealed that members of the family can be grouped in two clades, with P4H4 and P4H6 clustering together in clade II (**Supplemental Figure 2A**). P4H proteins contain signal peptides for their targeting to the secretory pathway, a transmembrane domain and a Fe(II) 2-oxoglutarate (OG) dioxygenase domain followed by a C-terminal Stichodactyla toxin (ShKT)-like domain with conserved cysteines only for the clade II (**Supplemental Figure 2B**). Fe(II) OG dioxygenase domain contain binding sites for Fe(II) (cofactor) and 2-oxoglutarate (co-substrate) essentials for P4Hs function (Gorres and Raines, 2010).

### Role of P4H4 and P4H6 in pollen germination and pollen tube growth

To evaluate the importance of prolyl hydroxylation during pollen germination and pollen tube growth, an analysis of Arabidopsis loss of function *p4h4* and *p4h6* mutants was performed. To this end, two T-DNA insertional mutant lines for *P4H4 (p4h4-1* and *p4h4-2*) and for *P4H6 (p4h6-1* and *p4h6-2*) were obtained from the Arabidopsis Biological Resource Center (ABRC) (**Figure 1A**) and a *p4h4-1 p4h6-1* double mutant was obtained by crossing. Pollen RT-PCR analysis confirmed down-regulation of *P4H6* transcript expression levels, but not *P4H4*, for each homozygous line (**Supplemental Figure 3**).

**Figure 1.**
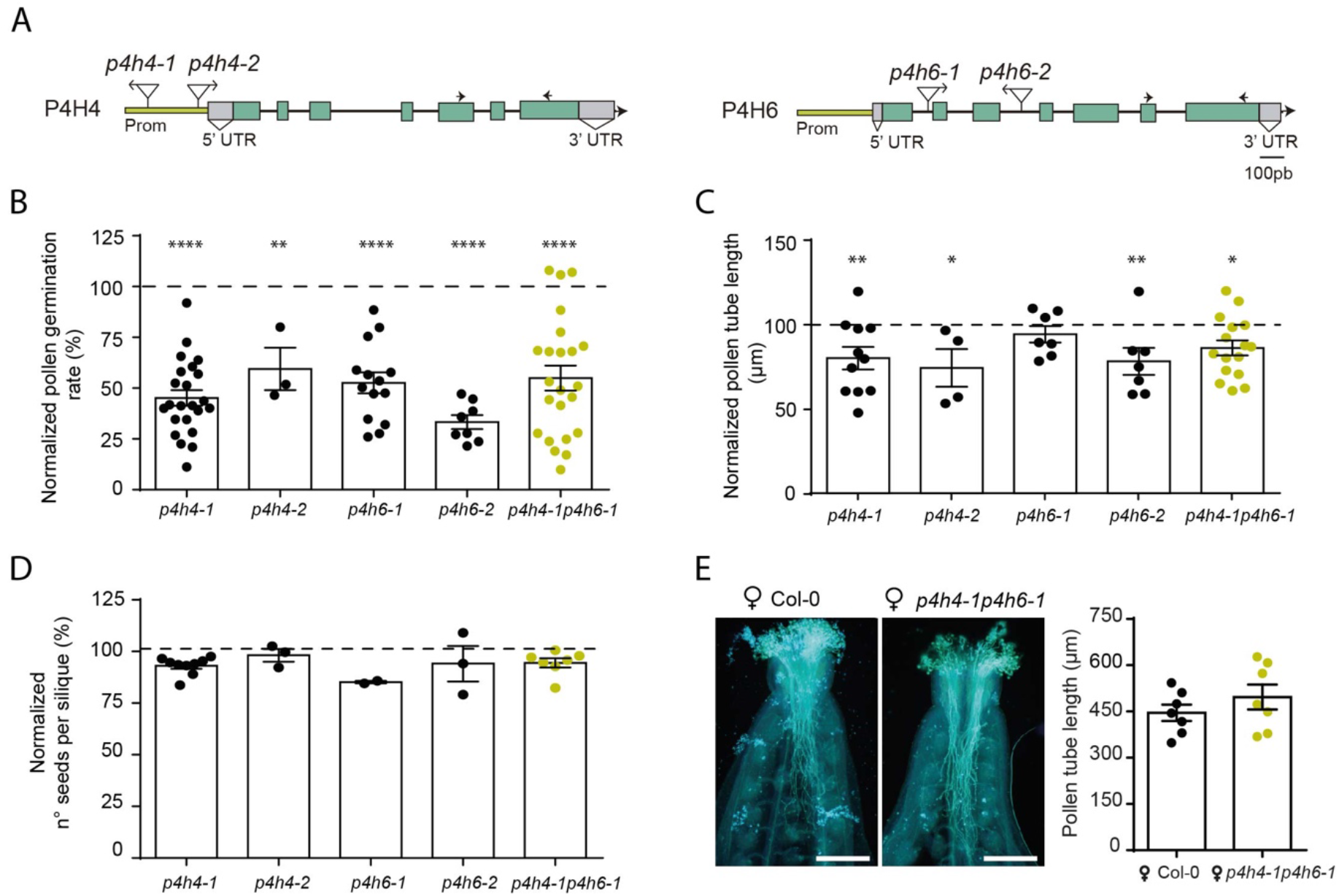
Analysis of Arabidopsis T-DNA loss of function *P4H4* and *P4H6* mutants. **A)** Structure of *P4H4* and *P4H6* genes. T-DNA insertional sites for each mutant line used in this study and primer hybridization regions for RT-PCR are indicated. Promoters (yellow), exons (green boxes) and introns (black lines) are also indicated. **B)** Pollen germination rate after 3 h of incubation *in vitro\**. **C)** Pollen tube length after 3 h of incubation *in vitro\**. **D)** Number of seed per silique (left panel)*. **E)** Representative images of WT Col-0 pistils hand-pollinated with Col-0 and *p4h4-1p4h6-1* pollen and stained with aniline blue 4 h after pollination (right panel); scale: 200 μm. Quantification of the pollen tube length within the pistil in *in vivo* assays (right panel); data is shown as the mean ± SEM and each symbol corresponds to a pollinated pistil. No significant differences were observed according to the Student’s *t* test. *Data is shown as the mean ± SEM and mean values were normalized to Col-0 that was set at 100%. Asterisks indicate significant differences with Col-0 according to a one-way ANOVA test: (*) p≤0.05, (**) p≤0.01 and (****) p≤0.0001.

Pollen germination ability of the loss of function mutants was evaluated *in vitro*. **Figure 1B** shows that after 3 h of *in vitro* incubation, a strong reduction in pollen germination was observed in *p4hs* mutant lines when compared to WT pollen (*p4h4-1*=45,9%, *p4h4-2*=59,5%, *p4h6-1*=52,6%, *p4h6-2*=33,3% and *p4h4-1p4h6-1*=54,9%%, where WT=100%). A statistically significant reduction in pollen tube length was also observed for most of the *p4h* mutants (**Figure 1C**). However, pollen of *p4h4-1 p4h6-1* double mutant did not display an additive phenotype compared to *p4h4-1* and *p4h6-1* single mutants (**Figure 1B-C**).

Despite the drastic reduction in pollen germination observed *in vitro*, a normal segregation ratio (2:26) was observed in the crosses between the two *p4h4-1 p4h6-1* homozygous single lines and the number of seeds per silique of all mutants was not reduced when compared to the WT (**Figure 1D**). Aniline blue stain of hand-pollinated pistils revealed that double mutant pollen tubes elongate properly within the pistil (**Figure 1E**). Finally, no significant differences in the content of pectin were observed in the cell wall of *p4h4-1p4h6-1* pollen tubes when compared to WT tubes (**Supplemental Figure 4**). All these results demonstrate that although *in vitro* pollen germination is highly reduced in the pollen *p4h* mutants, *in vivo* the pollen tubes grow normally and fertilize the ovules.

To test if the reduced germination rate observed in vitro in the *p4h4-1* mutant line corresponded to a bona fide phenotype, we decided to transform mutants with a complementation construct. To this end, transgenic lines expressing *P4H4* fused to *YFP* under its endogenous promoter and in the *p4h4-1* and *p4h4-1p4h6-1* homozygous mutant backgrounds were obtained. As shown in **Figure 2A**, the pollen germination phenotype observed in the *p4h4-1* line was complemented, confirming that although *P4H4* RNA levels were normal in the *p4h4-1* line, this mutant is indeed a loss of function mutant.

**Figure 2.**
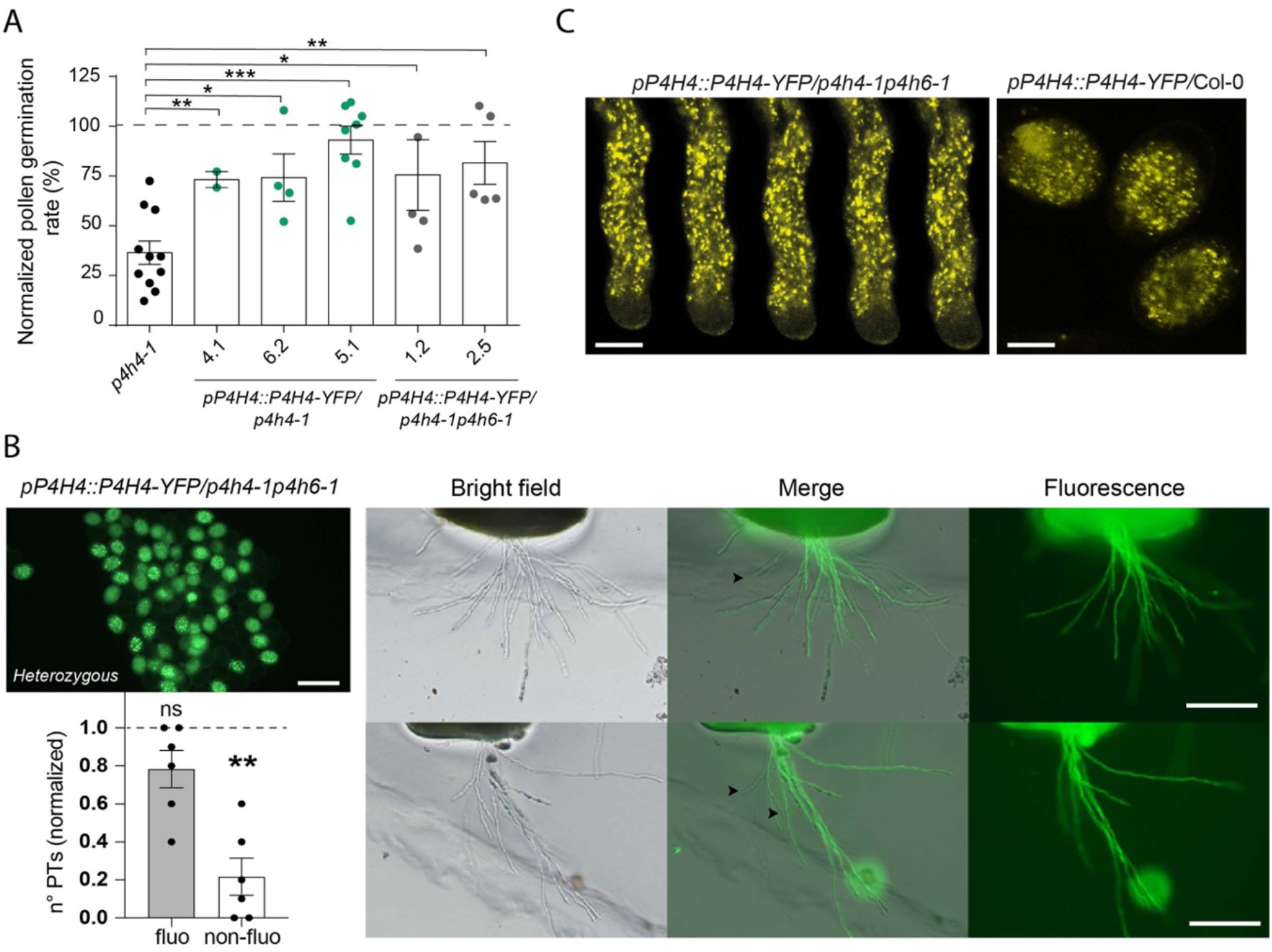
Complementation assay and subcellular localization of P4H4. **A)** Independent transgenic lines expressing *pP4H4::P4H4-YFP* construct in *p4h4-1* and *p4h4-1p4h6-1* mutant backgrounds. Pollen germination rate was calculated. Data is shown as the mean ± SEM and mean values were normalized to Col-0 that was set at 100%. Asterisks indicate significant differences with *p4h4-1* according to the Mann-Whitney non-parametric test: (*) p≤0.05, (**) p≤0.01 and (***) p≤0.001. **B)** Semi-*in vivo* pollen competition assay. Left, upper panel: Image of pollen from heterozygous complemented *pP4H4::P4H4-YFP/p4h4-1p4h6-1* transgenic line used in this experiment; scale: 50 μm. Two representative images of Col-0 pistils in a semi-*in vivo* assay. Black arrowheads indicate non fluorescent pollen tubes (P4H4-YFP^-^/*p4h4-1p4h6-1*); scale: 130 μm. Quantification and statistical analysis of six hand-pollinated pistils (left, bottom panel). Data is shown as the mean ± SEM of the percentage of fluorescent and non-fluorescent tubes normalized to Col-0 that was set at 1. Asterisks indicate significant differences with Col-0 according to a Mann-Whitney non-parametric test: (**) p≤0.01. **C)** Subcellular localization of P4H4 in a growing pollen tube and pollen grains. Confocal images of transgenic pollen expressing *pP4H4::P4H4-YFP* in Col-0 and *p4h4-1p4h6-1* backgrounds show that P4H4-YFP is localized in punctuate structures that resemble to Golgi apparatus/Golgi outgoing transport vesicles. Scale: 10 μm.

To compare pollen tube growth between the single mutant *p4h6-1* and the doble mutant *p4h4-1 p4h6-1*, we used the heterozygous transgenic line expressing *pP4H4::P4H4-YFP* in the *p4h4-1 p4h6-1* double mutant background in a semi-*in vivo* competition assay. The fluorescent half of the pollen grains (P4H4-YFP^+^/*p4h4-1 p4h6-1*) acts as a *p4h6-1* single mutant while the non-fluorescent half (P4H4-YFP^-^/*p4h4-1 p4h6-1*) lacks both *P4H4* and *P4H6* genes. Although there is a similar number of fluorescent and non-fluorescent pollen grains (**Figure 2B** left, upper panel), a higher percentage (75,4%) of fluorescent pollen tubes emerge from the cut style, suggesting that double mutant pollen (P4H4-YFP^-^/*p4h4-1 p4h6- 1*) has a reduced fitness to grow in the style compared to the complemented fluorescent pollen (P4H4-YFP^+^/*p4h4-1 p4h6-1*) (**Figure 2B**).

We also studied the subcellular localization of P4H4-YFP in pollen and pollen tubes. Other members of the P4Hs family, P4H2, P4H5 and P4H13, were previously shown to localize to the secretory endomembrane system in root hairs (Velasquez et al., 2015b). A similar localization pattern was observed for P4H4-YFP in pollen, as it is associated to punctuate structures that do not invade the clear zone of growing pollen tubes **(Figure 2C)**, resembling Golgi apparatus or/and Golgi outgoing transport vesicles (Tan et al., 2016; Jia et al., 2018).

### Chemical inhibition of P4Hs activity

Inhibition of P4Hs enzymatic activity was performed by applying the specific P4H inhibitors DP (α,α-dipyridyl) and EDHB (ethyl-3,4-dihydroxybenzoate) to the pollen germination media. The inhibitor DP chelates Fe^2+^, a necessary cofactor for P4Hs function, while EDHB competes with the co-substrate 2-oxoglutarate for the same binding site at the OG dioxygenase domain of P4Hs (**Supplemental Figure 2B**). **Figure 3A** shows that both inhibitors reduce WT pollen germination, as was shown for *p4hs* loss-of-function mutant lines (**Figure 1B**). Furthermore, at higher concentrations of both inhibitors, pollen tubes displayed an increase in the percentage of tubes showing abnormal morphologies and early bursting suggesting that hydroxylation of structural proteins is necessary for a proper cell wall assembly and pollen tube elongation. The half maximal inhibitory concentration (IC50) was calculated according to the pollen germination rate (IC50_DP=_ 15 μM and IC50_EDHB=_ 7 μM) (**Supplemental Figure 5**) and used in further experiments. Pollen germination of *p4h4- 1p4h6-1* double mutant shows a tendency to decrease after inhibitors treatment suggesting that other/s P4Hs might also be involved during pollen germination (**Figure 3B**).

**Figure 3.**
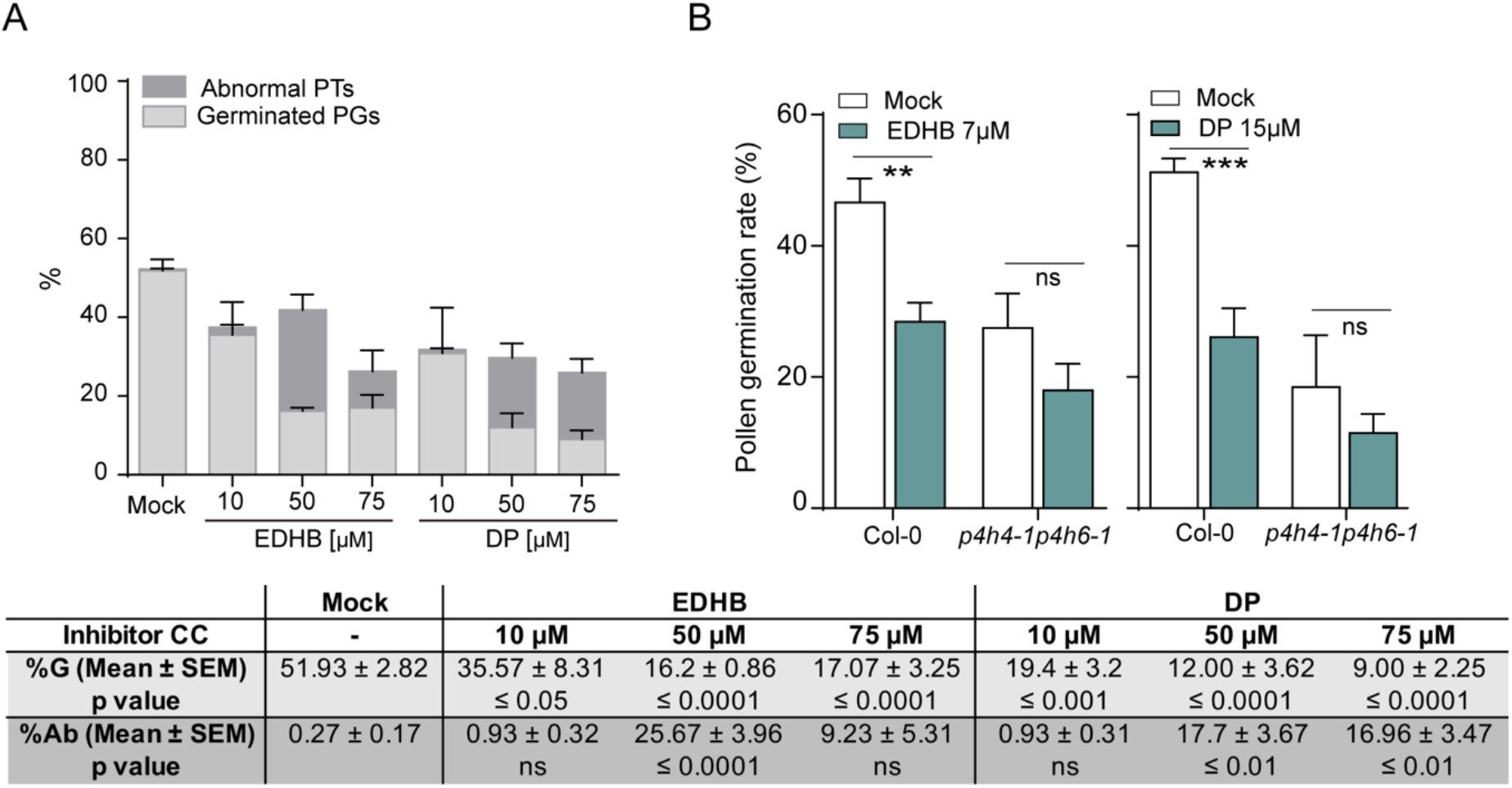
Chemical inhibition of P4Hs activity. **A)** Pharmacological inhibition of P4Hs activity in Col-0 pollen leads to a decrease in the pollen germination rate and an increase of abnormal morphologies of pollen tubes. Pollen germination rate was calculated after 3 h of *in vitro* incubation with or without (mock) the addition of specific P4H inhibitors, DP (α,α-dipyridyl) and EDHB (ethyl-3,4- dihydroxybenzoate) to the pollen germination media. Data is shown as the mean ± SEM for n=3 independent replicates with >200 pollen grains analyzed per replicate. Statistical analysis was performed according to a one-way ANOVA test (Table; bottom panel). %G refers to “percentage of germination” and %Ab to “abnormal pollen tubes”. **B)** Inhibitors treatment of Col-0 and *p4h4- 1p4h6-1* double mutant pollen with IC50 concentrations: IC50_EDHB=_ 7 μM and IC50_DP=_ 15 μM. Data is shown as the mean ± SEM of n≥6 independent replicates for each genotype per treatment. Asterisks indicate significant differences according to a one-way ANOVA test: (**) p≤0.01 and (***) p≤0.001.

### Leucine-rich repeat extensin LRX11 mislocalizes in *p4h4-1p4h6-1* pollen tubes

It was reported that LRXs are exported to the apoplast in polarized growing cells and localize mainly in the tip of pollen tubes (Ringli 2010; Ndinyanka Fabrice *et al*., 2018, Wang *et al*., 2018). In order to analyze whether prolyl-hydroxylation activity is necessary for proper localization of LRXs in the pollen tube cell wall, we explored the subcellular localization of *pLRX11::LRX11-GFP* in the *p4h4-1 p4h6-1* double mutant background. GFP fluorescence intensity was recorded along the pollen tubes by tracing a longitudinal line from the tip to the distal region (**Figure 4A**). **Figure 4B** shows that while 82.8% (± SEM=6.26) of the maximum intensity values of LRX11-GFP was recorded at the pollen tube tips in *Irx11* background, only 24.1% (± SEM=5.44) was detected in *p4h4-1p4h6-1* lines indicating that the final localization of LRX11 would depend on the presence of P4Hs. These results suggest that glycosylation of LRX11 mediated by P4H regulates the proper localization of LRXs in the apoplast of the pollen tube tip.

**Figure 4.**
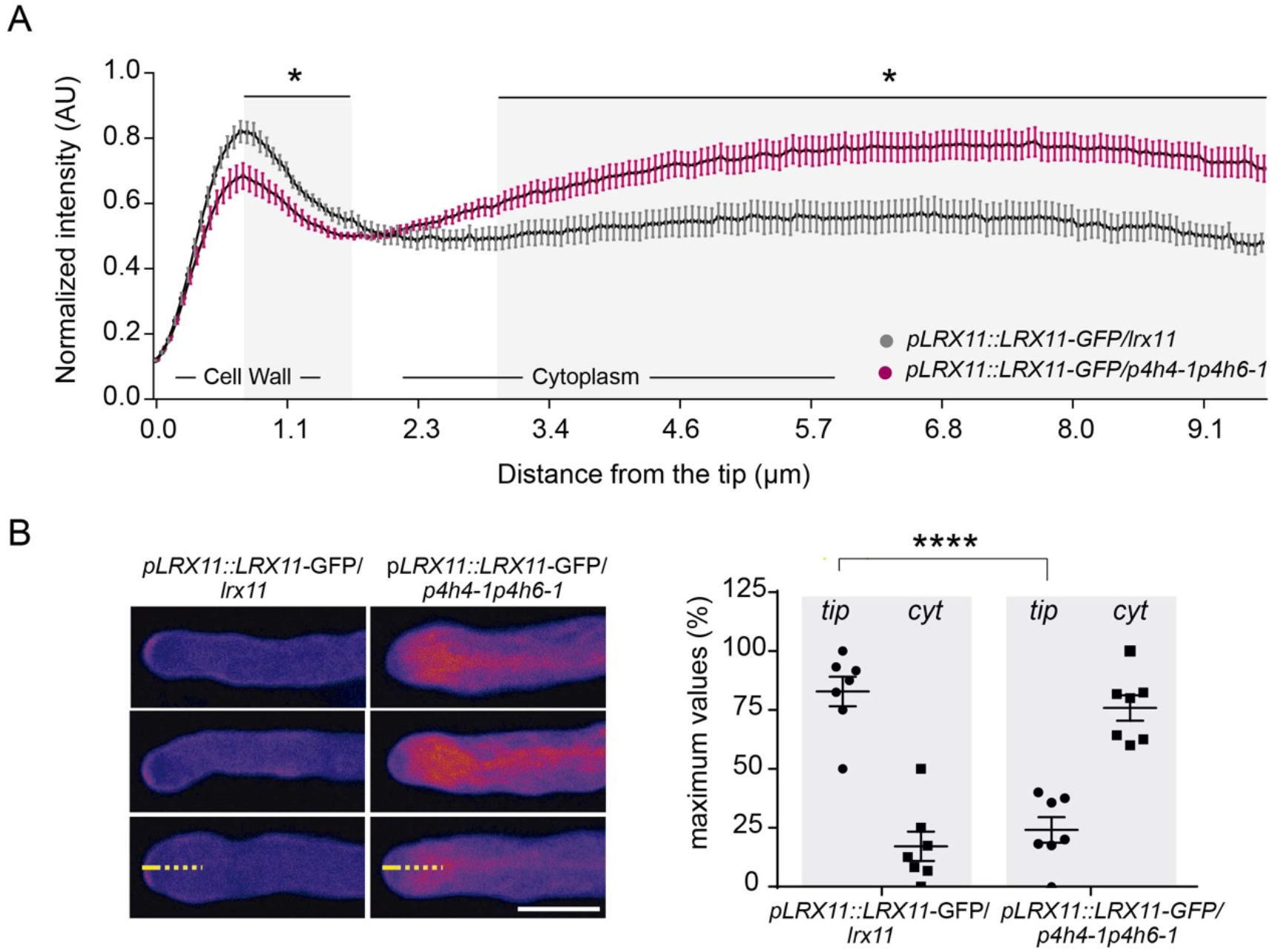
Analysis of LRX11-GFP localization in *lrx11* and *p4h4-1p4h6-1* mutant backgrounds. **A)** Measurements of LRX11-GFP intensity along the pollen tube by tracing a longitudinal line from the tip to the cytoplasm revealed that LRX11 mislocalizes from the cell wall in *p4h4-1p4h6-1* background. AU: arbitrary units. Data is shown as the mean ± SEM of n=7 independent experiments with ≥10 pollen tubes intensity measurements per transgenic line per experiment. Asterisks indicate significant differences according to a Multiple *t*-test; desired false discovery rate (Q) value was set to 5%. **B)** Confocal representative images of transgenic pollen tubes expressing the *pLRX11::LRX11- GFP* construct in *lrx11* and *p4h4-1p4h6-1* mutant backgrounds (left panel). Scale: 10 μm. Quantification of GFP intensity maximum values recorded at the tip (yellow line) and at the cytoplasm (cyt, dash yellow line) of transgenic pollen tubes (right panel); asterisks indicate significant differences according to Student’s *t* test: (****) p≤0.0001.

For a detailed analysis, co-localization assays of LRX11-GFP and the cell wall stained with PI were performed in plasmolyzed and non-plasmolyzed pollen tubes. In plasmolyzed *pLRX11::LRX11-GFP/lrx11* transgenic pollen tubes, LRX11-GFP co-localized with PI at the apoplast that corresponds to the pollen tube cell wall (**Figure 5A**). On the contrary, low LRX11-GFP signal was observed in the apoplast of *pLRX11::LRX11-GFP/p4h4-1p4h6-1* pollen tubes and no co-localization with PI was detected (**Figure 5A**). Then, the LRX11-GFP signal in the apoplast of plasmolyzed pollen tubes was quantified using the “Secretion Index” (SI; Beuder et al., 2020) as the ratio between the signal of cytoplasm/apoplast on a selected ROI. In *p4h4-1 p4h6-1* background, the calculated SI was 1.6 ± SEM=0.29 indicating a poorer localization of LRX11 in the cell wall when compared to the *pLRX11::LRX11-GFP/lrx11* control (SI = 0,66 ± SEM=0.09) (**Figure 5B**).

**Figure 5.**
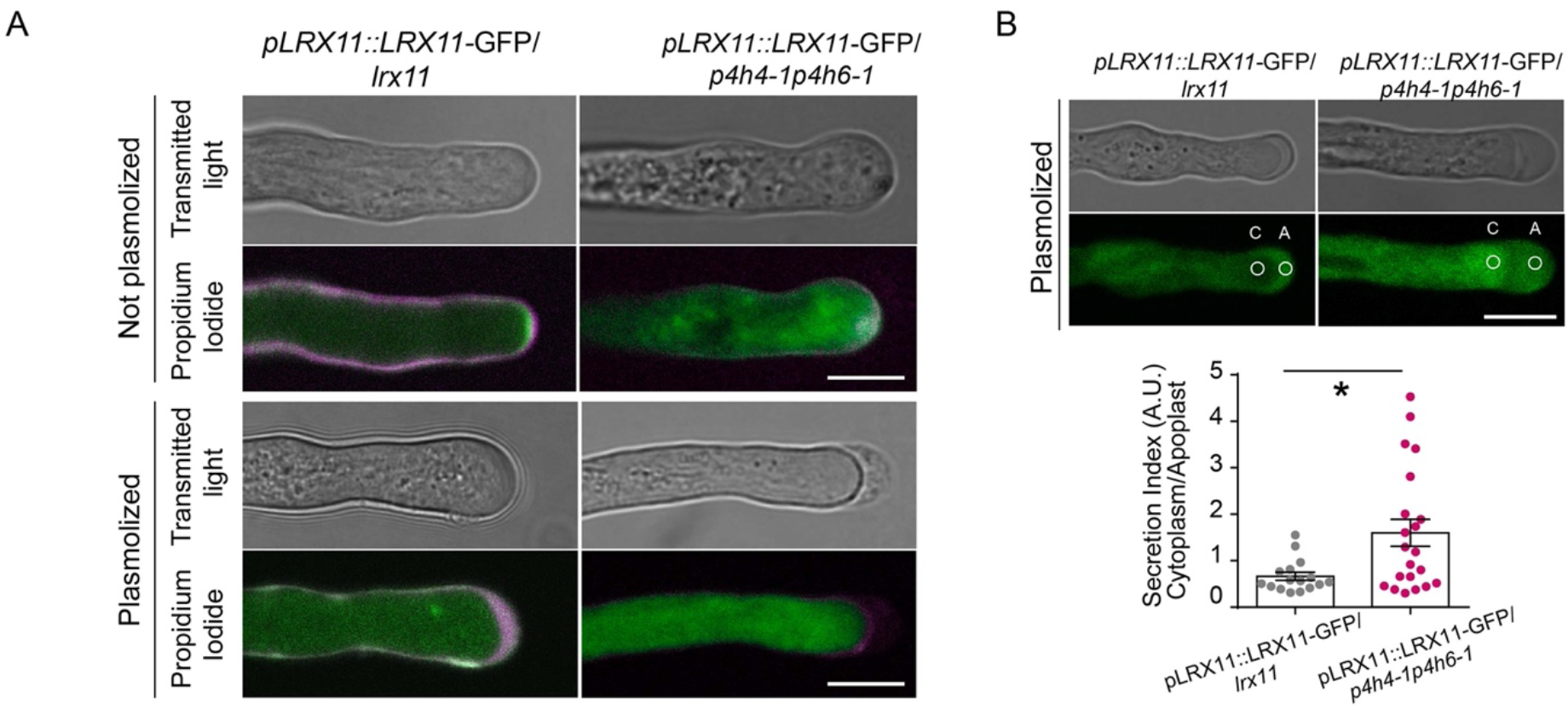
Analysis of LRX11-GFP localization in plasmolyzed pollen tubes. **A)** Merge confocal images of transgenic *pLRX11::LRX11-GFP* pollen tubes stained with propidium iodide for cell wall detection. Plasmolysis of pollen tubes was performed by applying mannitol 40% to the pollen germination media. Co-localization is shown in white. Scale: 10 μm. **B)** Quantification of LRX11-GFP intensity at the apoplast of plasmolyzed pollen tubes. Secretion index was calculated as the ratio of the GFP signal recorded in defined ROIs at the cytoplasm (C) and the apoplast (A). Scale: 10 μm. Data is shown as the mean ± SEM and asterisks indicate significant differences according to a Mann-Whitney non-parametric test: (*) p≤0.05. AU: arbitrary units.

The hydroxylation of prolines in glycopeptides is followed by a first step of *O*-glycosylation catalyzed by the glycosyltransferases HPAT1-3 which transfer an arabinose unit onto the Hyp (Ogawa-Ohnishi et al., 2013). In order to determine whether *O*-glycosylation by HPATs is occurring on LRXs, we analyzed the subcellular localization of LRX8-GFP and LRX11-GFP in the *lrx8+/-* or *lrx11+/-* mutant backgrounds respectively, and in the *hpat1hpat3* double homozygous mutant background. **Supplemental Figure 6** shows no differences in LRX11- GFP and a slightly difference in LRX8-GFP localization in *hpat1 hpat3* lines when compared to the respective *lrx8+/-* or *lrx11+/-* lines. This suggests that proline hydroxylation is sufficient for the proper localization of LRXs at the cell wall or else the action of still unknown glycosyltransferases involved in the *O*-glycosylation of the EXT domain of LRXs.

### Analysis of the hydroxylation profile over the EXT domain of LRX11

Western blot of pollen proteins from transgenic lines *pLRX8::LRX8-GFP, pLRX10::LRX10-GFP* and *pLRX11::LRX11-GFP* was performed using an anti-GFP antibody (**Figure 6A**). LRX8, LRX10 and LRX11 show two bands with a higher molecular weight than expected (102.8 kDa, 78.4 kDa and 75.2 kDa, respectively), suggesting the presence of post-translational modifications. Interestingly, when LRX11-GFP was immunoprecipitated in the *p4h4-1p4h6- 1* background, a single band of intermediate molecular weight was detected (**Figure 6B**).

**Figure 6.**
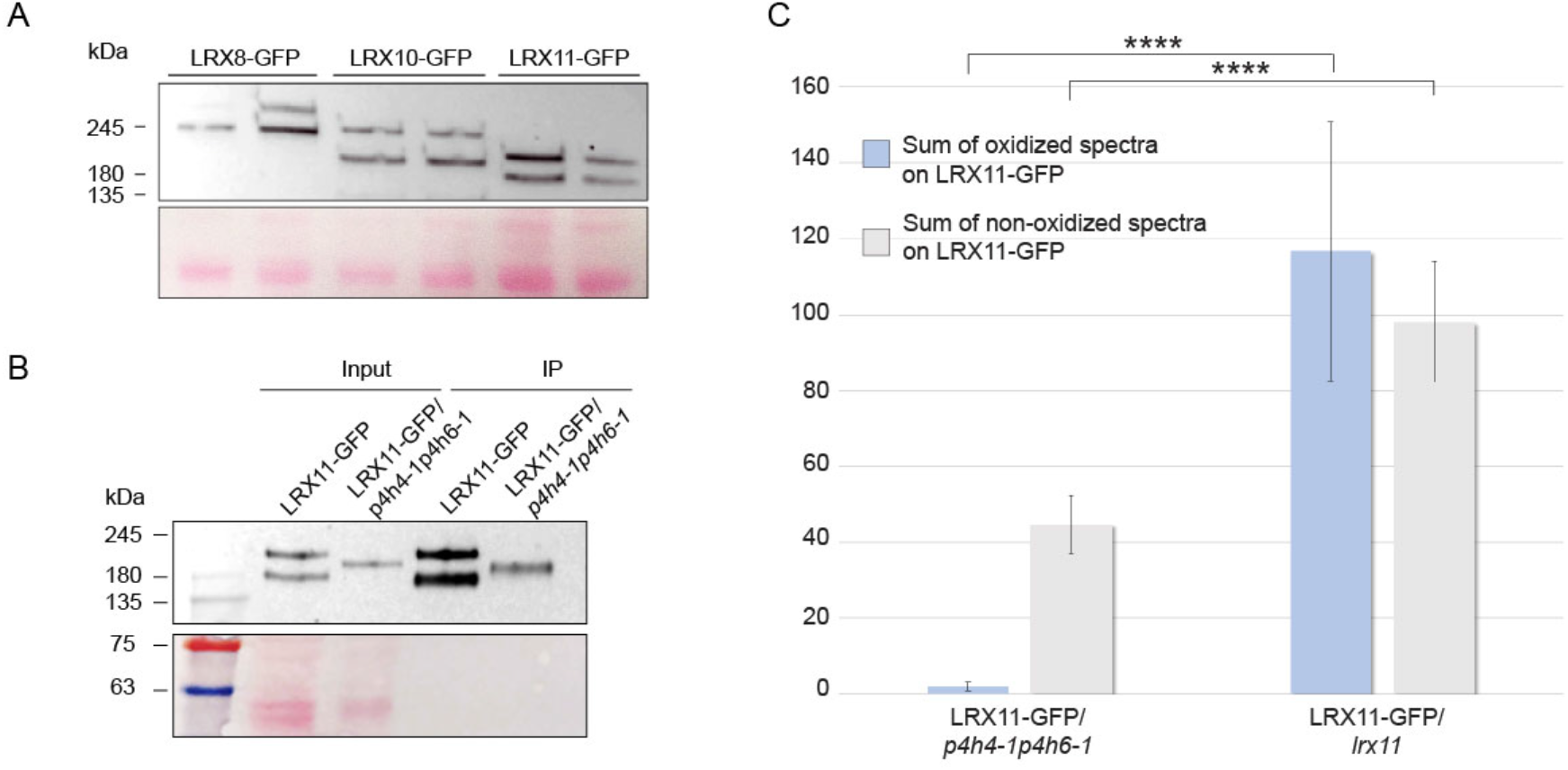
Western blot and LC/MS-MS analysis of pollen LRXs. **A)** Pollen LRX8/10/11-GFP display a higher molecular weight than expected (102.8 kDa. for LRX8, 78.4 kDa. for LRX10 and 75.2 kDa. for LRX11). Protein samples were obtained from pollen of *pLRX8::LRX8-GFP, pLRX10::LRX10-GFP* and *pLRX11::LRX11* -GFP transgenic lines and western blot was performed using an anti-GFP antibody; two biological replicates were included for each LRX. Ponceau S-stained nitrocellulose membrane is shown. **B)** Immunoprecipitation (IP) of LRX11-GFP using GFP-trap agarose beads from *pLRX11::LRX11-GFP/lrx11* and *pLRX11::LRX11-GFP/p4h4-1p4h6-1* protein samples. Ponceau S-stained nitrocellulose membrane is shown. **C)** Proline oxidation on LRX11-GFP in *p4h4-1 p4h6-1* and *lrx11* mutant backgrounds by LC/MS-MS analysis. Data is shown as the mean ± SD on three replicates; asterisks indicate, according to a t test, significant differences between oxidized and non-oxidized samples: (****) p<0.0001.

To investigate whether the absence of P4H4 and P4H6 affect the normal hydroxylation of LRX11, we performed an MS-MS analysis of GFP-immunoprecipitated pollen proteins from the *pLRX11::LRX11-GFP*/*lrx11* and *pLRX11::LRX11-GFP*/*p4h4-1p4h6-1* lines. Since trypsin hydrolysis sites on the proline rich C-terminal domain of LRX11 are not abundant, a double digestion with chymotrypsin and trypsin was performed to obtain shorter peptides. **Figure 6C** shows a statistically significant higher number of proline-oxidized LRX11 peptides in *pLRX11::LRX11-GFP/lrx11* pollen (117 peptides) when compared to *pLRX11::LRX11-GFP/p4h4-1 p4h6-1* (3 peptides) (se also **Supplemental Figure 7**). All these results suggest that P4H4 and P4H6 are responsible for the correct hydroxylation and proper folding of pollen LRX11.

## Discussion

LRX and other HRGP superfamily proteins require specific peptidyl-proline residues to be converted to trans-4-Hyp by P4H enzymes for proper cross-linking in the cell wall (Koski et al., 2007, 2009). Here, we show that pollen-specific P4H4/P4H6-catalyzed hydroxylation is required for pollen germination. Furthermore, chemical inhibition of P4Hs also affects pollen germination and an increase in the concentration of these inhibitors causes pollen tube bursting possibly due to alterations in cell wall structure.

In epidermal root cells, P4H5 is the main P4H that initiates and continues the Pro hydroxylation of EXTs, while P4H2 and P4H13 terminate hydroxylation at contiguous Pro residues (Velazquez et al. 2011; Velasquez et al. 2015b). The fact that P4Hs progressively hydroxylate Pro residues could explain the non-additive phenotype observed in the *p4h4-1 p4h6-1* double mutant. However, we show in semi-*in vivo* competition experiments that *p4h4-1 p4h6-1* pollen tubes displayed reduced fitness compared to *p4h6-1* single mutant. Despite this, fertilization is not compromised in the *p4h4-1 p4h6-1* mutant and *in vivo* analysis revealed that pollen tubes can elongate along the pistil.

P4Hs are membrane-bound enzymes that presumably localize to the Golgi apparatus where they form protein complexes and post-translational modifications occur (Yuasa et al., 2005; Velasquez et al. 2015b). Only when proline residues are hydroxylated as Hyp in the secretory pathway, they can be *O*-glycosylated with up to five Ara residues. The subcellular localization of P4H4 observed in pollen tubes would indicate its localization in Golgi but further analysis using specific organelle markers are required to support this observation.

*O*-glycosylation on EXTs results from hydroxylation of prolines and the addition of *O*-glycans in a sequential manner. The first arabinofuranose (Ara*f*) residue is transferred by HPAT1- HPAT3 (Ogawa-Ohnishi et al., 2013), the second by RRA1-3 (Reduced Residual Arabinose 1–3) (Egelund et al., 2007; Velasquez et al., 2011), the third Ara*f* by XEG113 (Xyloglucanase113) (Gille et al., 2009) and ExAD (Extensin Arabinose Deficient) transfers the fourth Ara*f* residue (Møller et al., 2017). The arabinosyltransferase that adds the fifth and final Ara unit has not yet been identified (Marzol et al. 2018). On the other hand, a single Serine-galactosyltransferase (SGT1/SerGT1) adds Gal to Ser in the repeated Ser-Hyp3-5 motifs (Saito et al., 2014). It has been described that *O*-glycans increase HRGP solubility, resistance to proteolytic degradation and thermal stability (Shpak et al., 2001; Kieliszewski et al., 2011; Lamport et al., 2011; Seifert 2020). Several Arabidopsis glycosyltransferases (GTs) mutants display root hair-defective growth phenotypes (Velasquez et al., 2011; 2015b), highlighting the role of *O*-glycans for the function of EXTs and related proteins during plant cell expansion. Moreover, the lack of Hyp-Ara in *hpat1* and *hpat3* loss of function mutants, causes an altered deposition of cell wall polymers in pollen tubes and leads to a reduction in pollen fertility (Beuder et al., 2020). Based on this, we propose a model in pollen in which the hydroxylation catalyzed by P4Hs on LRXs Ser-Pro(3-5) motifs is essential for their subsequent glycosylation (**Figure 7A**).

**Figure 7.**
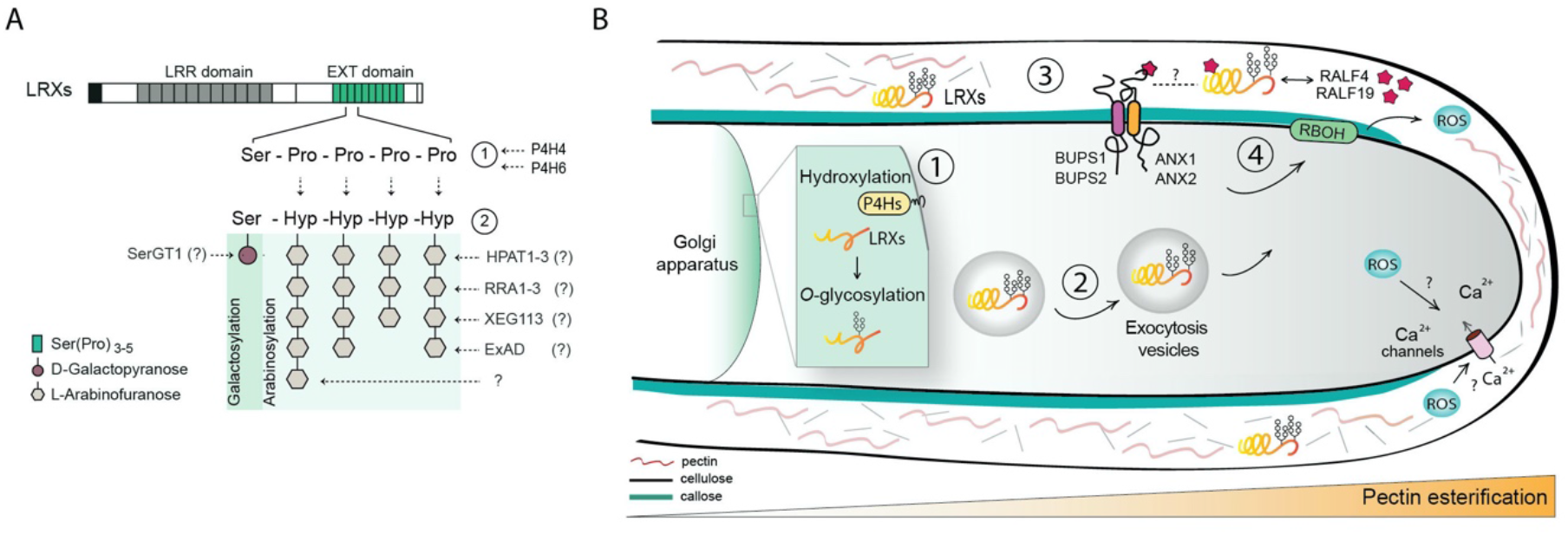
Proposed model: LRXs as targets of pollen P4Hs enzymes. **A)** Prolyl-4-hydroxylases, P4H4 and P4H6 catalyze the conversion of proline (Pro) to hydroxyproline (Hyp) originating new *O*- glycosilation sites (1). Different groups of glycosyltransferases are involved in a stepwise elongation of sugar chains over the Hyp and finally, a single galactose residue is added to serine (Ser) residues (2). SerGT1: serine-galactosyltransferase 1; HPAT1-3: Hydroxyprolil-*O*-arabinosyl-transferases 1-3; RRA1-3: Reduced Residual Arabinose 1-3; XEG113: Xyloglucanase 113; ExAD: Extensin Arabinose Deficient. **B)** P4H4 and P4H6 enzymes catalyze the hydroxylation of LRXs in the Golgi apparatus (1). Proper glycosylated LRXs are transported into exocytic vesicles (2) and delivered to the apoplast where participate in the crosslink and cell wall assembly (3). LRXs interact with RALF4/19 peptides in the cell wall activating a signaling pathway necessary (ROS and calcium) to sustain pollen tube polarized growth (4).

Glycosylated LRXs are delivered to the cell wall of polarized growing pollen tubes (**Figure 7B**) and are a linker between the plasma membrane and the cell wall (Fabrice et al., 2017, Wang et al., 2018). It was previously shown that the presence of Tyr cross-linking motifs in LRX1 is not required for its insolubilization in the cell wall (Ringli 2010) suggesting the existence of other robust interactions on the EXTs domain. Since pollen LRXs lack the Tyr motif, we explored whether Pro-hydroxylation and subsequent *O*-glycosylation are required for their retention in the cell wall. Thus, here we show that the correct hydroxylation at least in LRX11, is essential for its targeting. This result suggests that the lack of LRX11 proline hydroxylation, either genetically or pharmacologically, could alter its folding and its subsequent delivery to the apoplast, preventing an adequate anchorage in the cell wall and producing overaccumulation in the cytoplasm. This suggests that LRX11 is a target of P4H4 and P4H6 enzymes. On the other hand, arabinosylation would not be as important as hydroxylation, since pollen tube localization of LRX11 in double mutant *hpat1 hpat3* background was similar to WT.

LRXs play a main role in cell wall assembly, cell shape, and growth, but the role of each LRX in these processes remains unknown (Hall and Cannon, 2002; Cannon et al., 2008; Velasquez et al., 2011). There is high redundancy of LRXs among plant development. *lrx1/2* mutants show aberrant root hair morphologies (Baumberger et al., 2001; Baumberger et al., 2003a,b; Ringli, 2010) and the triple mutant *lrx3/4/5* shows defects in cell expansion in root cells (Draeger et al. 2015), possibly mediated by abnormal vacuolar expansion (Dünser et al. 2019). In addition, multiple mutants of pollen LRX8-11 show abnormal pollen tubes with an anomalous deposition of callose and pectin (Fabrice *et al*., 2017; Sede *et al*., 2018; Wang *et al*., 2018).

LRXs have been proposed to work together with receptor kinases of the CrRLK1L family (CrRLK1Ls) and Rapid Alkalinization Factors (RALF) peptides to monitor plant cell wall integrity status during polarized cell growth (Ge et al., 2017; Mecchia et al., 2017; Dunster et al., 2019; Herger et al., 2020; Somoza et al., 2021). **Figure 7B** shows a possible mechanism of action for pollen LRXs based on the interaction of LRX8 and LRX9 to RALF4 and RALF19 (Mecchia et al. 2017). RALF4/19 also bind to the extracellular domains of the pollen CrRLK1Ls receptor kinases such as ANX1/2 and BUPS1/2 (Ge et al. 2017). Similarly, the complex LRX3/4/5-RALF22/23-FERONIA coordinate growth under salt conditions (Zhao et al. 2018, 2020) and LRX1/5-RALF1-FERONIA in shoots and roots (Herger et al. 2020; Dünser et al. 2019).

Although the structural basis for the interaction between LRXs and RALFs peptides was recently established (Moussu et al. 2020), it remains unclear how *O*-glycans in the EXT domain of LRXs regulate these interactions. The EXT domain is variable among LRXs both in terms of length and motif (Borassi et al. 2016), suggesting that it has adapted to the specific cell wall architecture of numerous tissues as putative sensors of the cell wall integrity (Baumberger et al. 2003a,b; Marzol et al. 2018; Sede et al. 2018; Herger et al 2019). This may imply a more subtle effect of partial *O*-glycosylation on LRXs compared to completely non-hydroxylated/non-glycosylated LRXs.

## Experimental procedures

### Plant material and growth conditions

*Arabidopsis thaliana* ecotype Columbia plants (Col-0) were used as Wild-Type (WT) to perform all the experiments. Surface sterilization of seeds was performed by exposing them to chlorine fumes for 1.5 h. For stratification, sterilized seeds were incubated in petri dishes with 0.5X MS (Murashige and Skoog) basal medium at 4°C for 48 h in dark and then transferred to a growth chamber at 22°C under constant light. 10-day-old seedlings were transferred to soil and grown in a growth chamber under the same conditions.

### Genotyping of T-DNA mutant lines

Arabidopsis T-DNA mutant lines were obtained from the Arabidopsis Biological Resource Center (ABRC): for *P4H4* (AT5G18900) salk_102582 (*p4h4-1*) and salk_1229_D04 (*p4h4-2*) and for *P4H6* (AT3G28490), salk_067682 (*p4h6-1*) and salk_200619C (*p4h6-2*) were used. Genotyping of T-DNA mutant lines was performed by PCR reactions using genomic DNA extracted from rosette leaves. Primer for each mutant line were designed with *“iSect primer design tool”* available in signal.salk.edu/tdnaprimers.2.html (listed in **Table 1**). PCR reactions were performed as follows: 5 min at 94°C, 33 cycles of 30 sec at 94°C, 30 sec at 57°C/60°C and 1 min/Kb at 72°C with a final extension step of 10 min at 72°C.

**Table 1.**
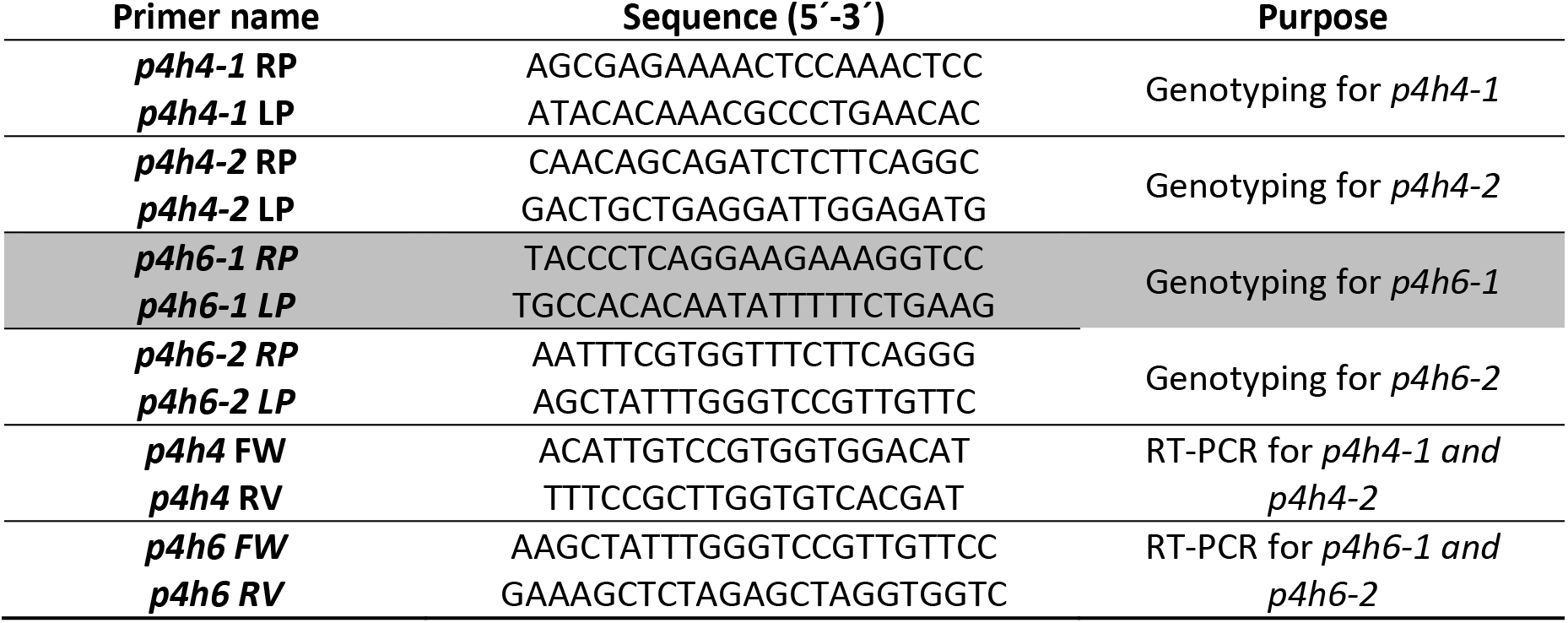
List of primers used in this work.

### Gene expression analysis

For RT-PCR analysis, one-day open flowers were collected and a total RNA isolation from mature pollen grains was performed with the TRIzol method according to the manufacturer’s instructions (Molecular Research Center, Inc). To achieve cDNA synthesis, equal RNA concentrations of each sample were used and incubated with reverse transcriptase MM-LV (*Promega*). A final PCR reaction using cDNA as template was performed as follows: 5 min at 94°C, 30 cycles of 30 sec at 94°C, 30 sec at 57°C/60°C, and 1 min/Kb at 72°C with a final extension step of 10 min at 72°C. Actin was used as an internal standard. All primers were designed to hybridize near to the C-terminal of each gene and are listed in **Table 1**.

### *In vitro* pollen germination and statistical analysis

One-day open flowers were collected and incubated over a wet paper in an incubation chamber for 30 min to allow pollen hydration as detailed in Sede *et al*., 2020. Pollen grains from anthers were brushed onto fresh semi-solid pollen germination medium prepared according to McCormick and Boavida, 2007 and incubated in a chamber at 22°C for 3 h. Only experiments with ≥ 30% germination rate were included in the analyses. Images were captured by selecting random fields with an epifluorescence Olympus BX41 microscope. Pollen tube length and pollen germination rate measurements were carried out using the Fiji software. In this study, the mean of ≥ 30 pollen tube length measurements per sample per genotype is consider as n=1 and ≥ 3 independent samples for each genotype were analyzed. For pollen germination rate determination ≥ 100 pollen grains per sample per genotype were analyzed and separated into two categories: not germinated pollen grains and germinated pollen grains (when the emerging pollen tube is twice as large as the perimeter of the pollen grain). The mean of these measurements was considered as n=1.

### Inhibitors treatment

P4Hs specific inhibitors, DP (α,α-dipyridyl) and EDHB (ethyl-3,4-dihydroxybenzoate) were added to the pollen germination media at different concentrations (10 μM, 50 μM and 75 μM) and pollen grains were incubated for 3 h. For IC50_DP_ and IC50_EDHB_ determination, a final concentration of 1 μM, 5 μM, 10 μM, 20 μM, 25 μM, 50 μM, 75 μM for DP and 1 μM, 2.5 μM, 7.5 μM, 10 μM, 15 μM, 25 μM, 50 μM for EDHB were selected. Inhibition curves (log[inhibitor] vs. response) were built according to WT pollen germination using the GraphPad Prism 6 software.

### *In silico* analysis

A heatmap of Arabidopsis *P4H* transcript expression pattern was built according to the ATH1 microarray data from Affimetrix in Col-0 background using the Genevestigator software. Expression patterns were compared with Arabidopsis pollen RNAseq data available in Loraine *et al*., 2013. Complete amino acid sequences of P4H1-P4H13 were aligned using MEGA software version 10.1.7 (Kumar et al., 2018) and a phylogenetic tree was built according to the Neighbor-Joining method.

### *In vivo* experiments

Arabidopsis WT pistils were emasculated and then manually pollinated. For *in vivo* assays, pistils were collected 4 h post pollination, fixated with an acetic acid/EtOH (1:3) solution and callose staining procedure was performed as detailed in Mori et al., 2006. For semi-*in vivo* experiments, Col-0 pistils were hand pollinated and the upper region of the stigma was cut 30 min post pollination, gently deposited over semi-solid pollen germination media and incubated in a chamber for 2.5 h at 22°C under constant light. Pistil images were obtained with an epifluorescence Olympus BX41 microscope.

### Confocal microscopy and statistics analysis

Pollen grains were germinated in semi-solid medium for 2 h as detailed before. For plasmolysis experiments, mannitol 40% was added to the pollen germination media after 2 h of pollen germination and incubated for other 30 min before image acquisition. The “Secretion index” was calculated as the ratio of fluorescence intensity recorded from two selected ROIs, one in the cytoplasm and the other in the apoplast of the pollen tube as indicated in Beuder et al., 2020. For cell wall staining, propidium iodide (0.2 mg/ml working solution) was added to the pollen germination media and images were acquired after 5 min incubation. Quantification of PI fluorescence intensity was performed at the perimeter of the pollen tubes (detailed in Sede *et al*., 2020) using the Fiji software.

### Transgenic lines

Transgenic plants carrying the *pLRX8::LRX8-GFP* / *lrx8 lrx9 lrx10* and *pLRX11::LRX11-GFP* / *lrx8 lrx9 lrx11* constructs were gently provided by the group of Dr. Yi Guo from Hebei Normal University, China. Transgenic plants were manually crossed with the *p4h4-1 p4h6-1* double mutant to obtain the stable lines *pLRX8::LRX8-GFP* / *p4h4-1 p4h6-1* and *pLRX11::LRX11- GFP/p4h4-1 p4h6-1* after several rounds of segregation. The cDNA *pP4H4::P4H4* was synthesized by Genscript company (Piscataway, NJ; http://www.genscript.com) on a pENTR/D-TOPO vector and then recombined in a pHGY destination vector for expression as a C-terminal YFP fusion protein. Transgenic plants carrying the *pP4H4::P4H4-YFP* construct were obtained by *Agrobacterium tumefaciens-mediated* transformation, using the floral dip method, and grown in 0.5X MS plates supplemented with Hygromycin for selection.

### Protein extraction and Western blot

One-day open flowers were collected and rapidly put in liquid N2. Total pollen protein extraction was performed as detailed in Chang and Huang, 2017. Protein concentrations of each sample were measured by a standard Bradford assay. For SDS-PAGE assay, 100 μg of protein of each sample were mixed with 2.5X of cracking buffer and boiled at 95°C for 5 min. Protein electrophoresis was performed in a 12% polyacrylamide gel and then transferred to a nitrocellulose membrane at constant 100 V for 1.5 h. For the immunoblot, membrane was incubated with GFP primary antibody (1:1000 dilution) and horseradish peroxidase (HRP)-labeled secondary antibody (1:1000 dilution) for luminescence detection.

### Immunoprecipitation and LC/MS-MS analysis

The LRX-GFP pollen protein sample was immunoprecipitated using GFP-trap agarose beads (ChromoTek) following the manufacturer’s instructions. For LC/MS-MS analysis, an enzymatic digestion of the samples with trypsin and chymotrypsin was performed to optimize cuts in the proline rich region of LRX11. A total of 3 replicates per genotype were included in the analysis. Pollen protein extracts from WT Col-0 plants were used as control. Immunoprecipitation and LC/MS-MS analysis. The LRX-GFP pollen protein sample was immunoprecipitated using GFP trap agarose beads (ChromoTek) following the manufacturer’s instructions. For LC/MS-MS analysis, samples were enzymatically digested with trypsin and chymotrypsin to optimize cuts in the proline-rich region of LRX11. A total of 3 replicates per genotype were included in the analysis. Pollen protein extracts from WT Col-0 plants were used as controls.

## Acknowledgements

We thank Cora MacAlister for providing the double mutant *hpat1 hpat3* seeds, the NASC (Ohio State University) for providing T-DNA lines seed lines and the Plateforme Protéomique Strasbourg Esplanade (CNRS) for their support in performing the mass-spectrometry analysis. MR, JME and JPM are investigators of the National Research Council (CONICET) from Argentina. This work was supported by grants from ANPCyT (PICT 2017-0076, PICT 2018-0504 and PICT2020-0103 to JPM; PICT2017-0066 and PICT2019-0015 to JME) and from the University of Buenos Aires Grants (UBACyT) to JPM. In addition, this research was also funded by ANID – Programa Iniciativa Científica Milenio ICN17_022, NCN2021_010 and Fondo Nacional de Desarrollo Científico y Tecnológico [1200010] to JME.

---

**Supp. Figure 1.**
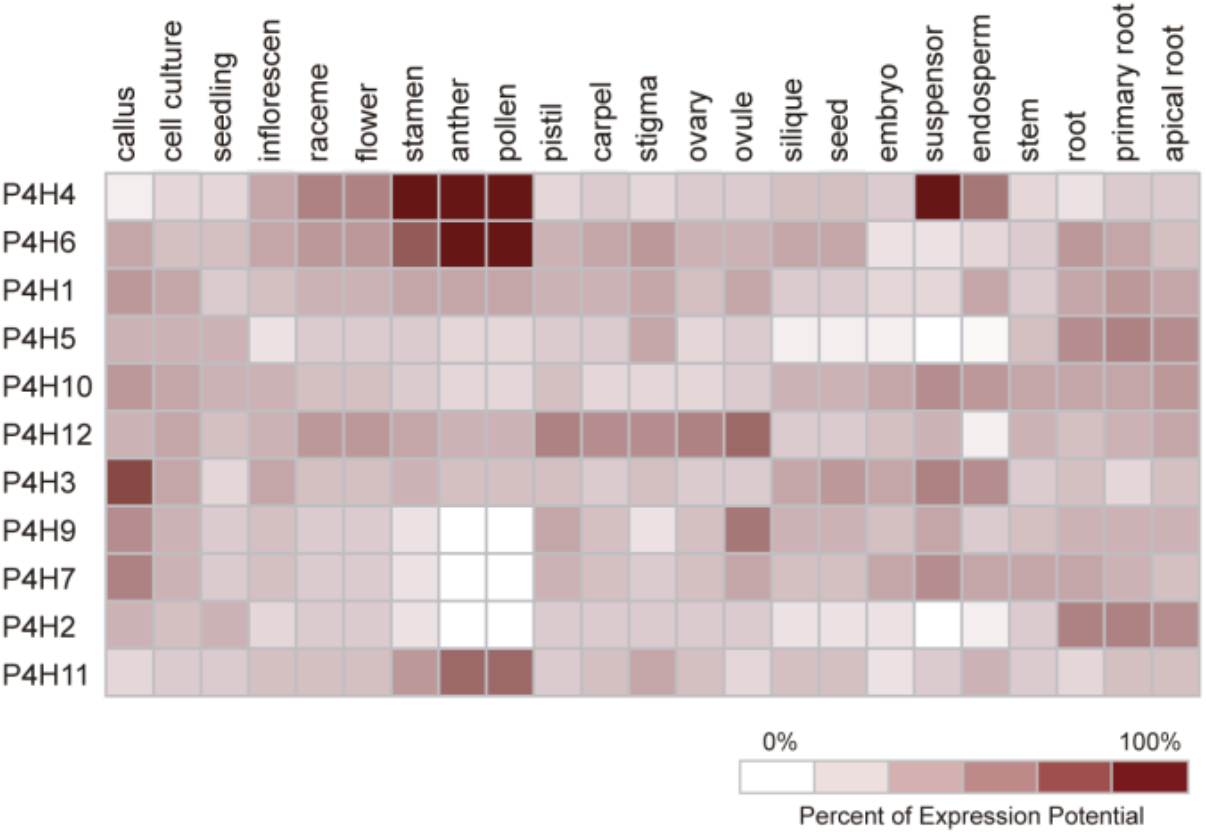
Heat map of *Arabidopsis thaliana P4Hs* gene expression profiles in mature pollen grains generated with Genevestigator according to ATH1 microarray data.

**Supp. Figure 2.**
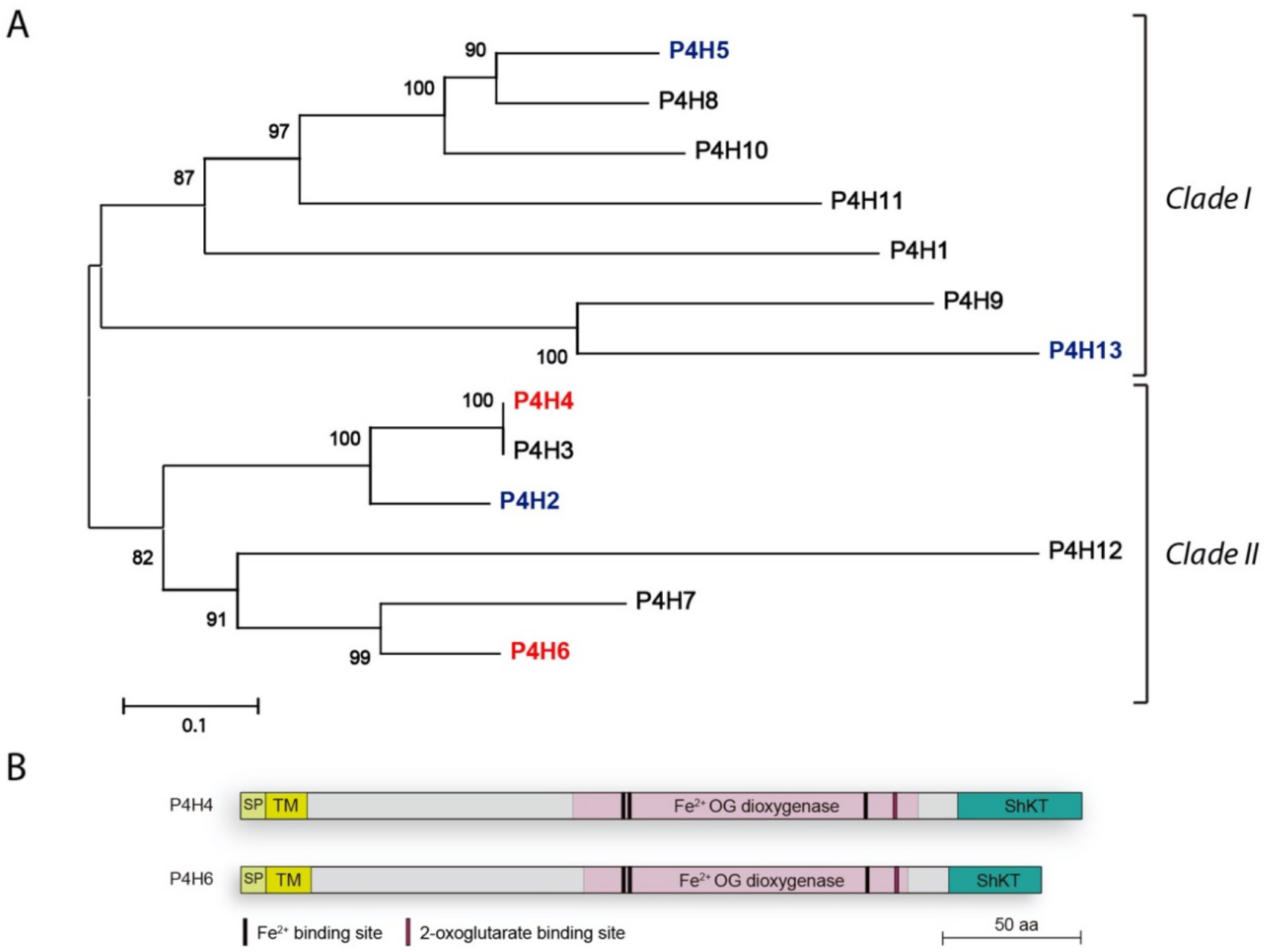
**A)** Phylogenetic tree of Arabidopsis P4Hs proteins built using the Neighbor-Joining method. Bootstrap values (1000 replicates) are shown next to the branches. Highly pollen-expressed P4Hs are indicated in red and P4Hs involved in polarized growth of root hairs are indicated in blue. Evolutionary analyses were conducted in MEGA-X software. **B)** Protein domain organization of P4H4 and P4H6. Signal peptide (SP), transmembrane domain (TM), 2-oxoglutarate (OG) dioxygenase domain and Cys-rich ShKT-like domain are indicated. Scale: 50 amino acids.

**Supp. Figure 3.**
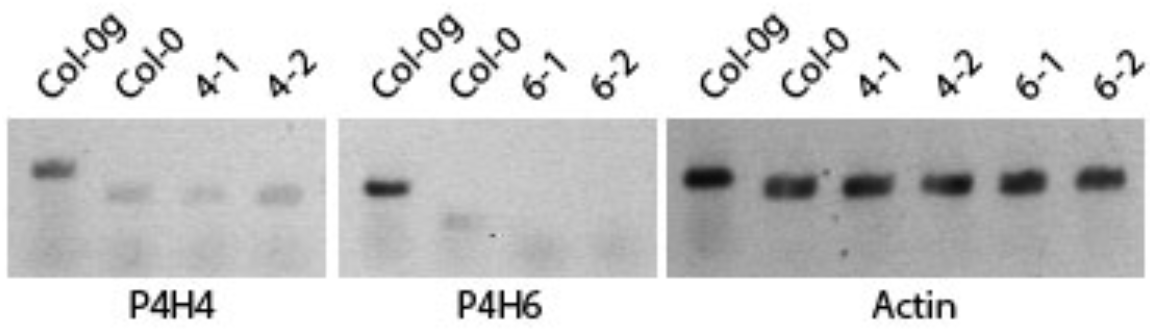
RT-PCR analysis of *P4H4* and *P4H6* transcript expression in pollen of Col-0 and T-DNA homozygous mutant lines *p4h4-1, p4h4-2, p4h6-1* and *p4h6-2*. Actin was used as a control. Genomic DNA (Col-0g) was included.

**Supp. Figure 4.**
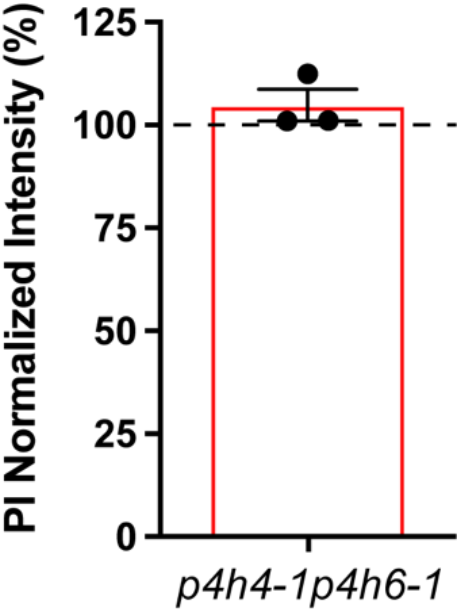
Quantification of pectin abundance in the cell wall of *p4h4-1p4h6-1* double mutant pollen tubes stained with propidium iodide. Measurements were performed at the perimeter of the pollen tube encompassing the tip and distal subapical region. Data is shown as the mean ± SEM of 3 independent experiments (n=3) and ≥12 pollen tubes were analyzed per experiment.

**Supp. Figure 5.**
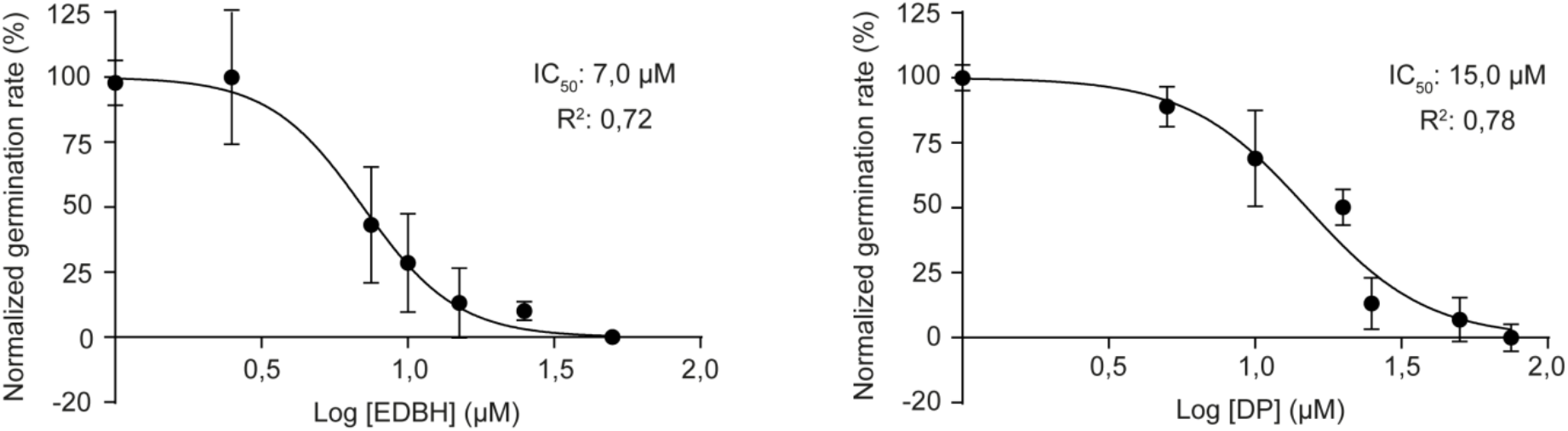
IC50 determination for P4Hs specific inhibitors. The half maximal inhibitory concentrations for DP (α,α-dipyridyl) and EDHB (ethyl-3,4-dihydroxybenzoate) were calculated according to the WT Col-0 pollen germination. Data is shown as the mean ± SEM of n≥3 replicates per treatment and ≥ 120 pollen grains analyzed per replicate.

**Supp. Figure 6.**
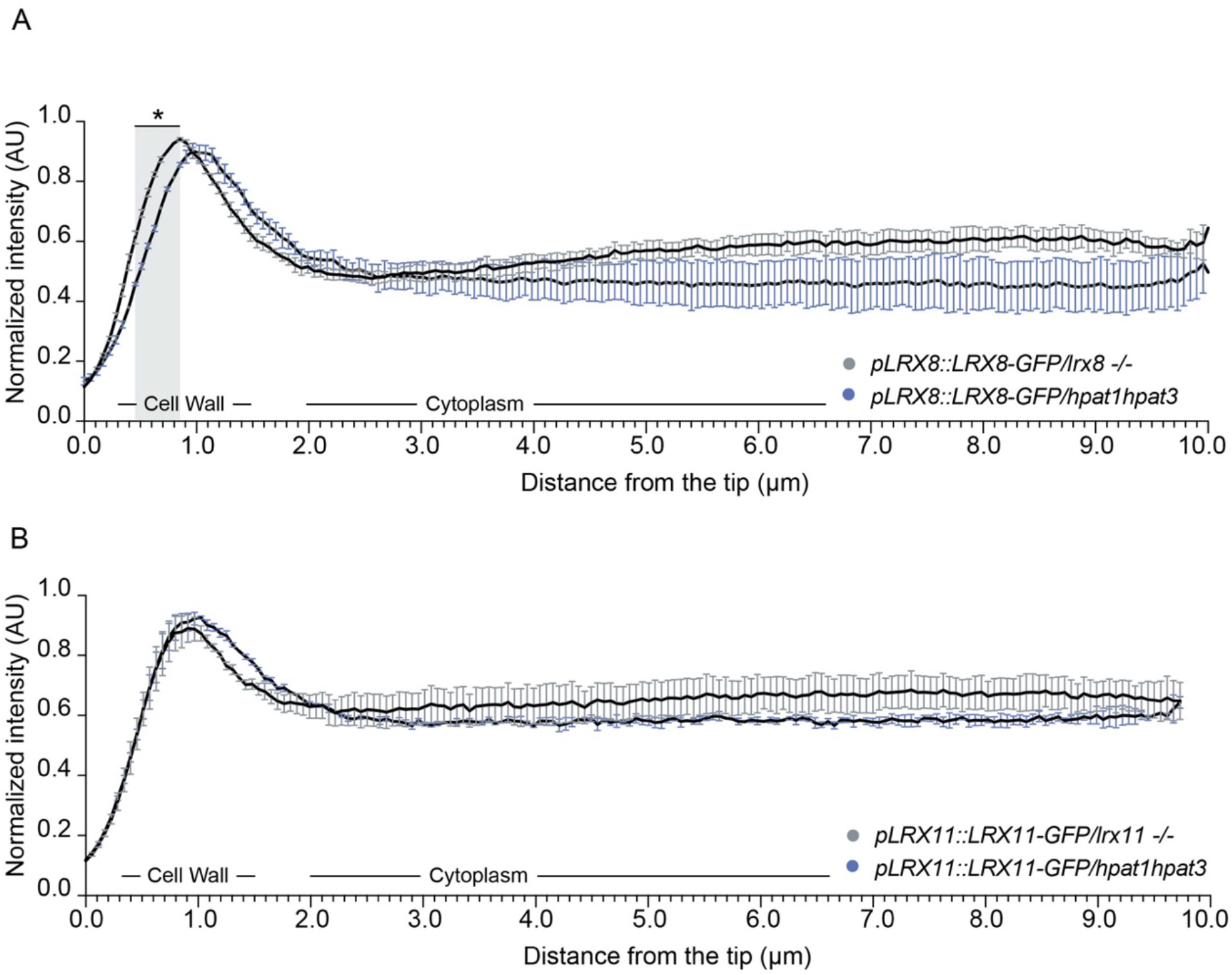
Analysis of LRX8-GFP and LRX11-GFP localization. **A)** Subcellular localization of LRX8- GFP in *lrx8* and *hpat1 hpat3* pollen tubes. **B)** Subcellular localization of LRX11-GFP in *lrx11* and *hpat1 hpat3* pollen tubes. Measurements of LRX8-GFP and LRX11-GFP intensity were performed along the pollen tube by tracing a longitudinal line from the tip to the cytoplasm. AU: arbitrary units. Data is shown as the mean ± SEM of n=3 independent experiments with ≥7 pollen tubes intensity measurements per transgenic line per experiment. Asterisks indicate significant differences according to a Multiple t-test; desired false discovery rate (Q) value was set to 5%.

**Supp. Figure 7.**
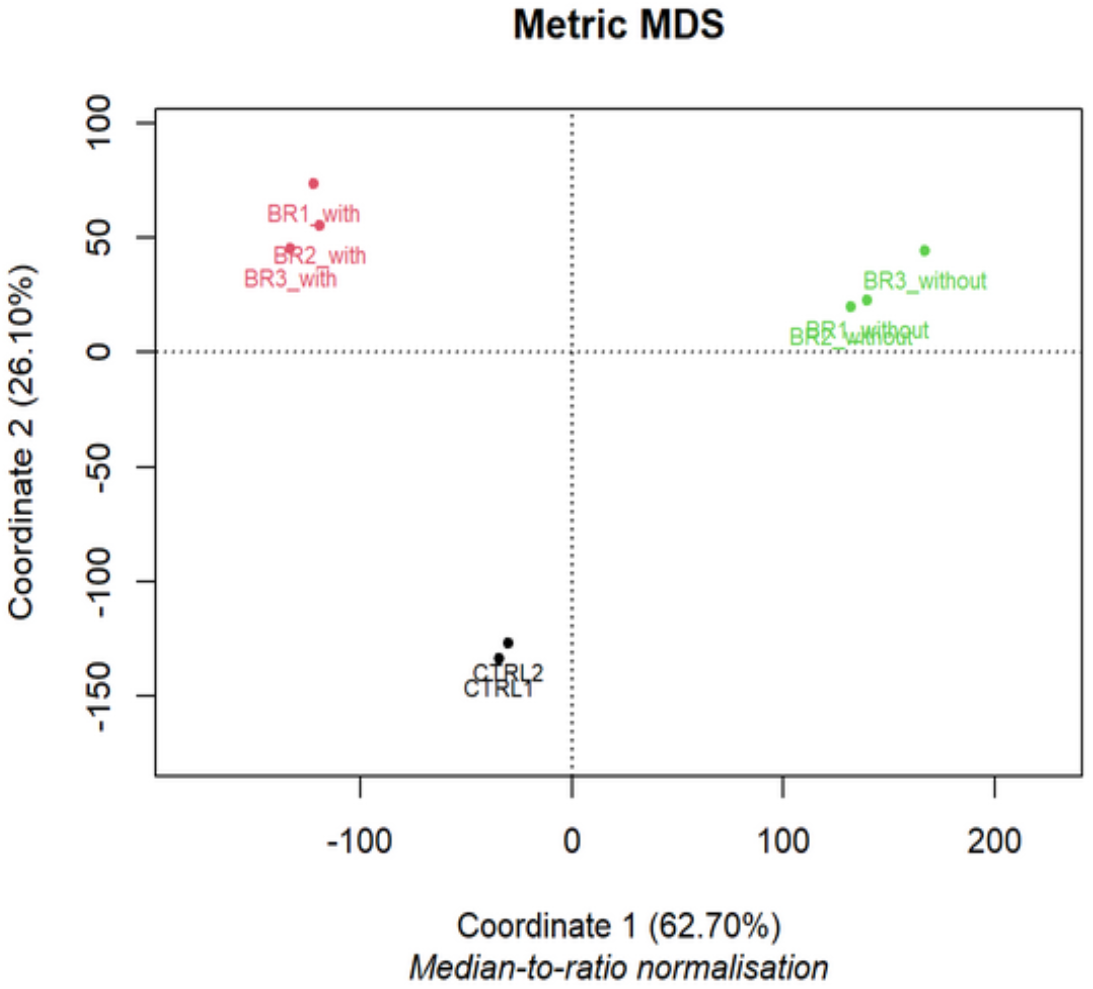
Multidimensional scaling plot of samples analyzed by LC/MS-MS. Three replicates for *pLRX11::LRX11-GFP/lrx11* (in red) and *pLRX11::LRX11-GFP/p4h4-1p4h6-1* (in green) and two replicates for the control (in black) are shown.

